# Top-down modulation of neural envelope tracking: the interplay with behavioral, self-report and neural measures of listening effort

**DOI:** 10.1101/815365

**Authors:** Lien Decruy, Damien Lesenfants, Jonas Vanthornhout, Tom Francart

## Abstract

When listening to natural speech, our neural activity tracks the speech envelope. Moreover, recent research has demonstrated that this neural envelope tracking can be affected by top-down processes. The present study was designed to examine if neural envelope tracking is modulated by the effort that a person expends during listening. Five measures were included to quantify listening effort: two behavioral measures based on a novel dual-task paradigm, a self-report effort measure and two neural measures related to neural phase synchronization and alpha power. Electroencephalography responses to sentences, presented at a wide range of subject-specific signal-to-noise ratios, were recorded in thirteen young, normal-hearing adults. A comparison of the five measures revealed different effects of listening effort as a function of speech understanding. Reaction times on the primary task and self-reported effort decreased with increasing speech understanding. In contrast, reaction times on the secondary task and alpha power showed a peak-shaped behavior with highest effort at intermediate speech understanding levels. We found a positive association between envelope tracking and speech understanding. While a significant effect of listening effort was found on theta-band envelope tracking, the effect size was negligible. Therefore, our results suggest that listening effort is not a confound when using envelope tracking to objectively measure speech understanding in young, normal-hearing adults.

## Introduction

Recent advances in signal processing have enabled the measurement of neural tracking of the low amplitude modulations of natural speech (Lalor & Foxe, 2010; Ding & Simon, 2014). As these modulations, also called the speech envelope, are essential for speech understanding (Shannon *et al*., 1995), it has been suggested to measure neural tracking of the speech envelope to assess speech understanding. Recent studies have shown an increase in envelope tracking with increasing speech understanding supporting its potential as an objective measure of speech understanding (Ding & Simon, 2012; Vanthornhout *et al*., 2018; Decruy *et al*., 2019; Etard & Reichenbach, 2019; Lesenfants *et al*., 2019). Nevertheless, it is important to take into account that top-down processes can influence neural envelope tracking. For example, there is evidence that prior knowledge affects envelope tracking. More specifically, Cervantes Constantino & Simon (2018) found that envelope tracking persists when removing segments of the speech. Di Liberto *et al*. (2018) showed enhanced envelope tracking when vocoded speech was made intelligible by first presenting the unprocessed speech. Research investigating the cocktail party problem has also demonstrated a substantial influence of attention. Neural envelope tracking of the attended, target speech was found to be enhanced compared to the ignored, competing speech (Ding & Simon, 2012; O’Sullivan *et al*., 2015; Das *et al*., 2016). In view of this, the question arises if the top-down process listening effort, defined as the deliberate allocation of mental resources to overcome obstacles in goal pursuit when carrying out a listening task (Pichora-Fuller *et al*., 2016), also modulates envelope tracking.

For several reasons there has been a large interest in the concept of listening effort. Firstly, it is common that two individuals achieve a similar performance on a clinical speech-in-noise test but experience a different degree of effortful listening in daily life situations (Anderson Gosselin & Gagné, 2010). Secondly, research in which performance is equalized for two groups shows that older normal-hearing (NH) participants expend significantly more effort during a speech-in-noise test than young NH adults (Anderson Gosselin & Gagné, 2011). To overcome these problems, new speech-in-noise tests could be developed that require an equal amount of effort. Current, more feasible approaches involve the inclusion of listening effort measures that provide additional information to current speech-in-noise tests. We have to note, however, that the concept of listening effort is still widely debated partly due to the existence of many different methods to measure it. In the next sections, a short literature overview is given.

To measure listening effort during a behavioral experiment, a single-task or a dual-task paradigm can be used. Single-task paradigms usually involve a speech-in-noise test in which the reaction time to the recalling of heard sentences, is registered (Gatehouse & Gordon, 1990; Houben *et al*., 2013; Pals *et al*., 2015). A dual-task paradigm consists of two tasks and is based on the assumption that a person has a limited amount of cognitive resources that can be allocated to the two tasks (Kahneman, 1973). If the primary task is difficult, a person will allocate all their resources to this task, resulting in a limited amount of resources remaining for the secondary task. Accordingly, the performance on this secondary task represents the effort expended on the primary task. Although dual-task paradigms are widely used, no consensus is yet achieved about which type of dual-task should be used (Gagné *et al*., 2017).

Next to behavioral tasks, self-report measures are often used to investigate listening effort since these can be easily implemented in an existing speech-in-noise test (Luts *et al*., 2010; Zekveld *et al*., 2010; Rudner *et al*., 2012; Wendt *et al*., 2014). In a study of Picou et al. (2011), participants were instructed to indicate their degree of perceived effort using a visual analog scale from 0 (no effort) to 10 (lots of effort). Although self-report measures are easily acquired, the outcome relies on the participant’s ability to reliably estimate his/her listening effort as well as the participant’s interpretation of the question.

To avoid this, physiological measures could be used since these do not require the cooperation of the participant. For example, Bernarding *et al*. (2014) proposed the phase synchronization of electroencephalography (EEG) responses to sentences as an objective measure of listening effort. More specifically, they found that less effortful conditions are reflected in the EEG by a uniform phase distribution whereas demanding conditions are associated with a clustered, non-uniform phase distribution (Bernarding *et al*., 2013, 2014). Next to phase, other EEG measures have also been associated with listening effort, e.g. alpha oscillations (Obleser *et al*., 2012; Miles *et al*., 2017; Dimitrijevic *et al*., 2019) or amplitude of event-related potentials (Bertoli & Bodmer, 2016). Lastly, several studies have also shown that the pupil size and eye movements can be used as physiological indicators of processing load (Zekveld *et al*., 2010, 2018; Koelewijn *et al*., 2014; Wendt *et al*., 2014). In spite of this, the neural mechanisms underlying the changes in pupil size are complex, resulting in different outcomes compared to other effort measures (Zekveld *et al*., 2018; Alhanbali *et al*., 2019). In addition, pupillometry is affected by several factors such as age, ambient light, and requires expensive equipment as well as specialized training which makes it difficult to implement it in the clinic or research (Houben *et al*., 2013; Winn *et al*., 2018).

Because listening effort can be measured using different methods, researchers often use a combination in their studies. For example, Wu *et al*. (2016) measured listening effort using two dual-tasks as well as a rating scale. They found that participants performed better on the easy secondary task than the hard secondary task, indicating that less effort was needed to complete the easy task. In contrast, the ratings demonstrated higher effort for the easy versus hard task. Different measures can thus lead to different outcomes and interpretations.

Recent reviews have addressed this issue and suggest that different measures tap into different aspects of listening effort (Lemke & Besser, 2016; Pichora-Fuller *et al*., 2016; Ohlenforst, Zekveld, & Jansma *et al*., 2017; Alhanbali *et al*., 2019). For this reason, we chose to include five measures in this study. The first two measures involve a dual-task paradigm in which we register (1) the reaction time to the recalling of heard sentences and (2) the reaction time to a secondary task. According to previous research (Picou & Ricketts, 2014) forcing competition between the same pool of resources creates a dual-task paradigm that is more sensitive to changes in listening effort. Hence, we created a new secondary task in which we ensured that the primary and secondary task required the allocation of similar cognitive resources. We chose to create a verbal working memory task as it plays an important role in speech perception (Rönnberg *et al*., 2013; Peelle, 2018). In addition to the dual-task, we included a (3) self-report measure in order to measure a different aspect of listening effort, perceived effort (Lemke & Besser, 2016). For two reasons, we decided to also measure the (4) phase synchronization and (5) the alpha power of neural responses to sentences. Firstly, studies have shown an association between changes in these measures and changes in listening effort (alpha: Obleser *et al*., 2012; phase: Bernarding *et al*., 2014). Secondly, phase synchronization and alpha power are, in contrast to the previous effort measures, direct measures of brain activity similar to neural envelope tracking (Peelle, 2018; Dimitrijevic *et al*., 2019).

In the present study, our first aim was to compare the five listening effort measures as a function of speech understanding. All measures were evaluated across a wide range of signal-to-noise ratios (SNRs) because recent dual-task and pupillometry studies have shown a non-linear, peak-shaped behavior of listening effort as a function of SNR (Zekveld *et al*., 2014; Wu *et al*., 2016; Ohlenforst, Zekveld, & Lunner *et al*., 2017). More specifically, it was shown that low SNRs not always resulted in extreme effort, but can also be associated with minimal effort when participants give up (Pichora-Fuller *et al*., 2016; Peelle, 2018). By evaluating each measure across a wide range of subject-specific SNRs, we extend the current understanding of listening effort measured by reaction times to heard sentences, phase synchronization and alpha power. We hypothesize that all measures, except self-reported effort, show a peak-shaped behavior across a wide range of SNRs since these measures reflect processing load, i.e. the amount of cognitive resources allocated to a task. For the self-report measure, we expect that perceived effort is highest at the low SNRs and decreases with increasing SNR.

The second aim of this study involves how much each listening effort measure accounts for the unexplained variability in envelope tracking beyond speech understanding. Previous studies have shown that envelope tracking has the potential to be used as an objective measure of speech understanding (Vanthornhout *et al*., 2018; Decruy *et al*., 2019; Lesenfants *et al*., 2019; Verschueren *et al*., 2019). However, young NH participants show differences in neural envelope tracking that can not be fully attributed to changes in SNR or speech understanding. In addition, recent studies have observed a lower envelope tracking for equally intelligible sentences presented without a masker compared to sentences presented at a high SNR (Das *et al*., 2018; Lesenfants *et al*., 2019). We hypothesize that these two unexplained differences in envelope tracking are related to changes in listening effort based on the findings of following behavioral studies. Wu *et al*. (2016) have found that listening effort, measured using a dual-task, is generally characterized by a peak-shaped behavior. When assessing the individual results, however, they found that the shape of the curve varied substantially across participants. In addition, Houben *et al*. (2013) have shown that participants achieve similar speech understanding for sentences presented without a masker compared to sentences presented at +4 dB SNR, but expended significantly less effort for the condition without a masker. Besides this, it is suggested that the delta and theta frequency band play different roles in the encoding of speech (Ding & Simon, 2014; Etard & Reichenbach, 2019; Lesenfants *et al*., 2019). Since the precise role remains debated, we also investigated if listening effort modulates neural envelope tracking for both frequency bands or only for the delta- or theta-band.

## Material and methods

### Participants

Thirteen Flemish speaking participants, aged between 17 and 28 years, participated in this study. All participants had normal hearing as their pure tone thresholds were better than 25 dB HL at all octave frequencies in both ears (125 Hz up to 8 kHz). During a short interview, the participant’s medical history and education were questioned because it is known that serious concussions, medication for the treatment of insomnia (Van Lier *et al*., 2004) and learning disabilities such as dyslexia (Power *et al*., 2016; De Vos *et al*., 2017) can affect brain responses. The hand and ear preference of the participants were also determined using a Flemish, modified version of the laterality preference inventory of Coren (1993). Lastly, a word reading test, cognitive screening test and working memory test were administered to assess reading and cognitive skills which are needed for completing the secondary task (see Dual-task paradigm). With regard to the word reading test (één-minuut test; Brus & Voeten, 1973), all participants obtained a score higher than percentile 10, indicating no persistent reading and/or spelling difficulties. With regard to the cognitive tests, all participants passed the cognitive screening test (> 25/30 on Montreal Cognitive Assessment; Nasreddine *et al*., 2005) and achieved a score higher than 27/60 on the working memory test (Flemish computerized version of Reading Span Test, RST; Van Den Noort *et al*., 2008; Vercammen *et al*., 2017). This study was approved by the Medical Ethics Committee UZ KU Leuven / Research (reference no. S57102 (Belg. Regnr: B322201422186) and S58970 (Belg. Regnr: B322201629016)). All participants took part voluntarily and gave their written informed consent.

### Behavioral experiment

After screening, participants first completed the behavioral experiment that took approximately two hours (overview figure 1). For all participants, this took place at the research group ExpORL of the KU Leuven, in a double-walled, soundproof booth or in a quiet room at the participant’s home. First, the subject-specific SNRs were determined. Second, the speech-in-noise test and verbal working memory test were administered simultaneously in a dual-task paradigm. In the following sections, the tests and two behavioral measures of listening effort are described in more detail.

**Figure 1:**
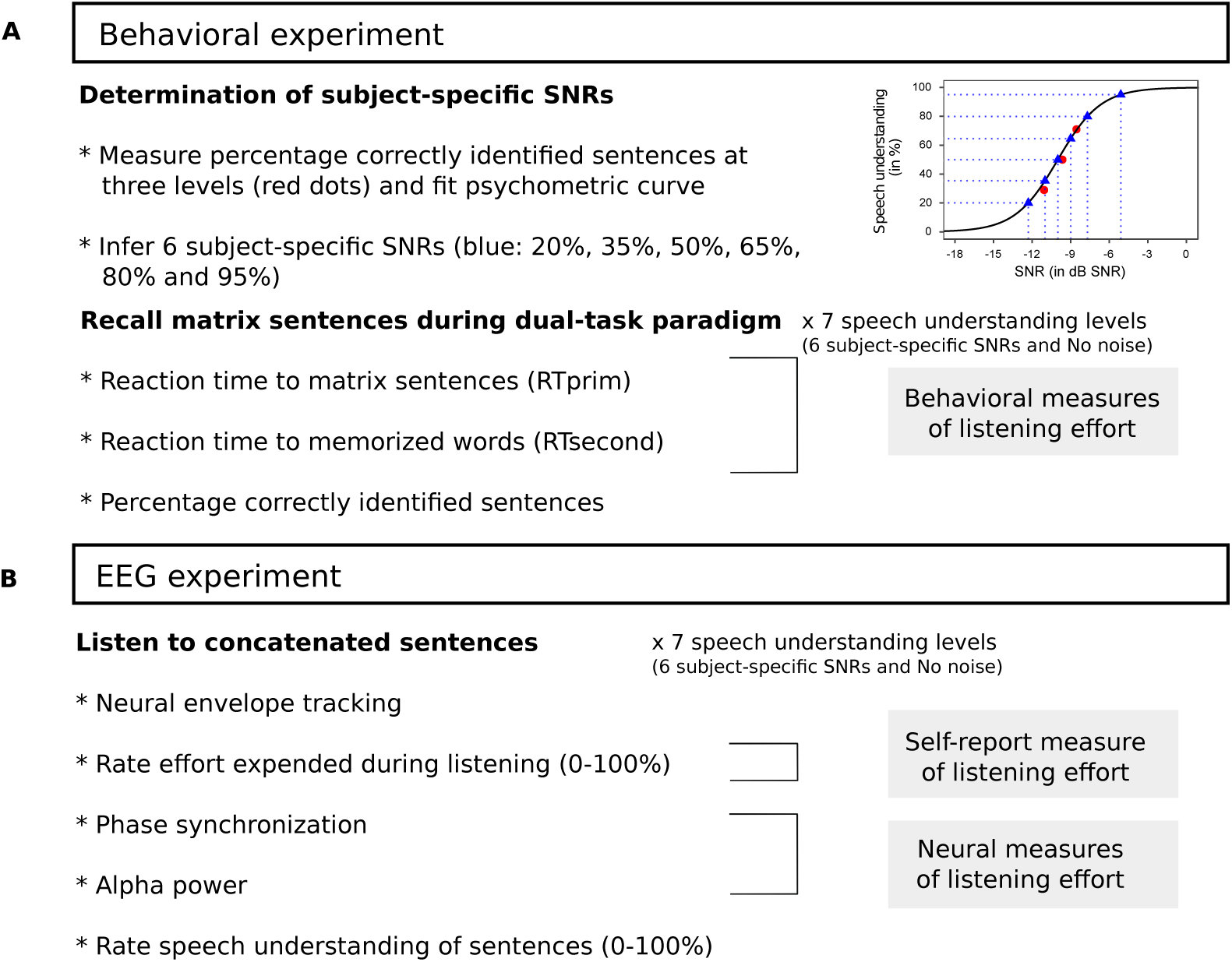
Overview of study design.

#### Determination of a wide range of subject-specific SNRs

In this study, we wanted to investigate listening effort across a wide range of SNRs because recent studies demonstrated a peak-shaped behavior of listening effort as a function of SNR (Zekveld & Kramer, 2014; Wu *et al*., 2016; Ohlenforst, Zekveld, & Lunner *et al*., 2017). To ensure this, six subject-specific SNRs were determined prior to the dual-task paradigm. The Flemish matrix sentence test (Luts *et al*., 2014) was used as speech material and consists of sentences with a fixed syntactical structure of 5 word categories “name, verb, numeral, color and object” (e.g. “Emma buys three blue bikes”). Participants were instructed to recall the sentences out loud using a 5 × 10 matrix containing 10 possibilities for each word of the sentence. We motivated participants to guess based on the matrix when it was more difficult to understand the sentences. When participants did not know the answer, they could also choose to give no answer. The percentage of correctly repeated words per list/SNR was calculated to obtain a speech understanding score.

To obtain the SNRs, the matrix sentences were masked with a stationary speech-weighted noise with the same long-term average spectrum as the matrix sentences. Three adaptive procedures (Brand & Kollmeier, 2002) converging to 29%, 50% and 71% speech understanding (SU) were administered to determine the six subject-specific SNRs. This was done by using the result of each adaptive procedure to fit a sigmoid function per participant using the following formula 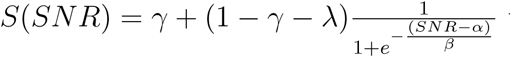 with *γ* the guess-rate fixed to zero, *λ* the lapse-rate fixed to zero, *α* the midpoint and *β* the slope. From this fit, we consequently inferred a subject-specific SNR per SU-level (20%, 35%, 50%, 65%, 80%, 95% SU). These subject-specific SNRs were used for both the behavioral and EEG experiment.

#### Dual-task paradigm

We created a new dual-task paradigm in which both tasks require the allocation of similar cognitive resources: 1) the primary task is a standardized speech-in-noise test and 2) the secondary task is a verbal working memory test, especially developed for this study.

During the dual-task paradigm, participants were instructed to prioritize the primary speech-in-noise task to ensure that the performance on the secondary task reflects the effort expended on the primary task (limited cognitive capacity theory of Kahneman, 1973). As illustrated in figure 2, a dual-task trial consisted of three screens. First, participants were instructed to read and remember three consonant-vowel-consonant (CVC) words (e.g. “cat, hut, pen”) which were presented on a computer screen for three seconds. Second, a blank screen appeared, a matrix sentence was played and participants were asked to recall the sentence out loud. Third, one of the three CVC words reappeared, e.g. “pen” and participants were instructed to push, as fast and accurate as possible, the button that corresponded with the original position at which the word was presented, e.g. “pen: third position”. Per trial, it was randomly chosen which word reappeared. Note that this secondary task also relies on short-term and visuospatial working memory since participants are instructed to remember the words and their position on the screen. Nevertheless, the explicit instruction to read the words, activates verbal working memory which is the same cognitive function that speech-in-noise tests rely on. In view of this, our test is likely to be more sensitive to changes in listening effort than other visual tasks used in dual-task paradigms (Picou & Ricketts, 2014; Gagné *et al*., 2017).

**Figure 2:**
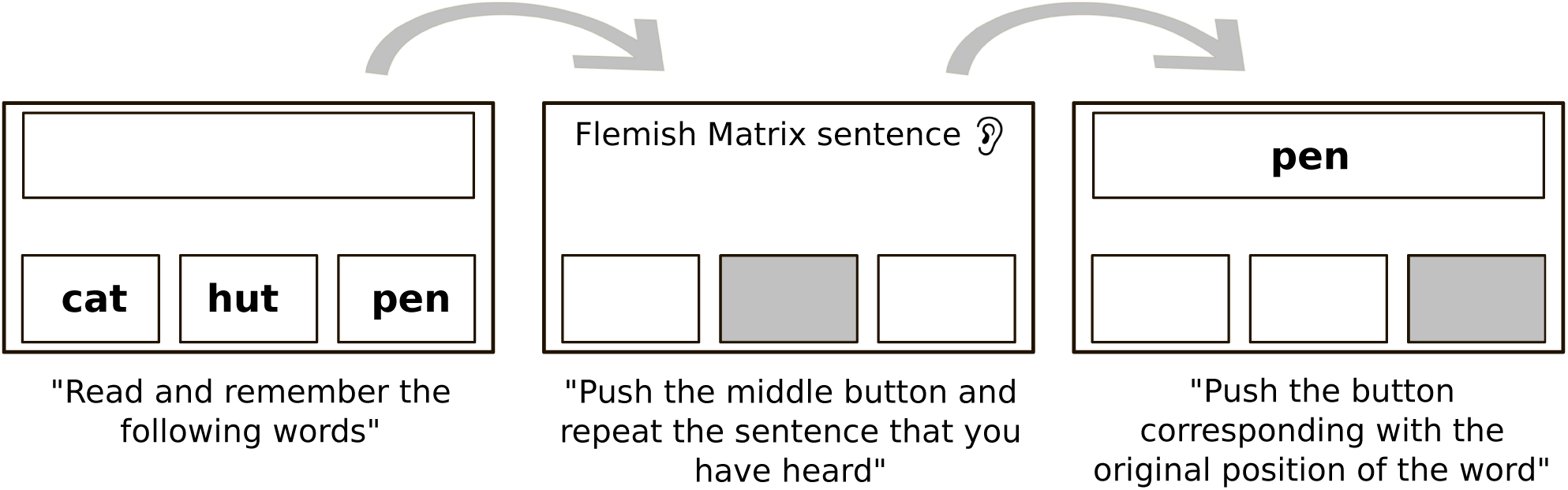
Overview of the dual-task paradigm. Per trial three screens were presented. First, three CVC words appear, then a matrix sentence is presented and finally, one of the three words reappears.

Since studies have shown that reaction times during single speech-in-noise tasks can be used as a behavioral correlate of listening effort (Gatehouse & Gordon, 1990; Pals *et al*., 2015), we asked the participants to push the middle button when recalling the matrix sentence (second screen). The reaction time (RTprim) was calculated between the end of the auditory presentation of the sentence and the recall of the participant (i.e. when the participant pushes the middle button). In dual-task studies, often the reaction time on the secondary task is used as a measure of listening effort (Anderson Gosselin & Gagné, 2010; Gagné *et al*., 2017). In this study, the reaction time (RTsecond) was calculated as the time between the reappearance of the CVC word and the answer of the participant (i.e. when the participant pushes the button corresponding with the original position). Similar to Houben *et al*. (2013), we included both correct and incorrect trials because we did not want to remove valuable information about the difficult secondary task conditions. To assess listening effort as a function of speech understanding, six matrix sentence lists of each 20 dual-task trials were administered at six fixed, subject-specific SNRs (20%, 35%, 50%, 65%, 80%, 95% SU) and one list was presented without a masker (No noise).

Prior to the dual-task paradigm, the speech-in-noise test and verbal working memory test were first administered separately in a single-task paradigm to exclude the bias due to procedural learning for the dual-task paradigm. Similarly to the dual-task paradigm, seven lists of the matrix sentence test were administered (six subject-specific SNRs and No noise). Participants also completed 10 runs of 10 trials of the verbal working memory test, because pilot results showed no further improvement in reaction times after 100 trials.

#### Reaction time preprocessing

Using reaction times as an outcome measure often requires several pre-processing steps because reaction times are not normally distributed but are always positive and skewed to the right. In this study, we pre-processed our reaction times in three steps. First, we calculated the median of the reaction times across trials, per SNR and per participant (Ratcliff, 1993). Second, reaction times shorter than 200 ms (Whelan, 2008) were removed as well as reaction times longer than 2.24 * (*MAD/*0.6745) + *M* with *MAD* defined as the median absolute deviation and *M* being the sample median (MAD-median rule: Wilcox *et al*., 2013). After outlier removal, we normalized the reaction times per participant in a last step by subtracting each value with the median for the “No noise” condition and dividing this by the median of the “No noise” condition (Houben *et al*., 2013; 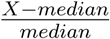; Gagné *et al*., 2017).

#### Auditory and visual stimulation

For the presentation of the matrix sentences, we used a laptop connected to a RME Hammerfall DSP Multiface II or RME Fireface UC soundcard (RME, Haimhausen, Germany). The sentences were presented using the software platform APEX (Dept. Neurosciences, KU Leuven) (Francart *et al*., 2008). For 8 of the 13 participants, the target speech stimuli and masker were presented diotically through ER-3A insert phones (Etymotic Research, Inc., IL, USA). For the remaining 5 participants, the stimuli were presented monaurally to the right ear. To create the subject-specific SNRs, the level of the noise was adjusted while the speech level was fixed at an intensity of 62 or 65 dB SPL (A weighted) for diotic and monaural stimulation respectively. Before administering the experiments, all stimuli were first calibrated with a type 2260 sound level pressure meter, a type 4189 half-inch microphone and a 2cc coupler (Bruel & Kjaer, Copenhagen, Denmark). Similar to matrix sentences for the primary task, the three CVC-words for the secondary task were presented using the software platform APEX. The reaction times during the verbal working memory task were registered using APEX in combination with three large buttons mounted on a wooden board. At an equal distance of each button, a hand was drawn on the wooden board to indicate the place at which participants could place their preference hand for pushing the buttons.

### EEG experiment

After the behavioral experiment, an EEG experiment of approximately two and a half hours was conducted to investigate the association between neural envelope tracking and listening effort (overview figure 1). During the EEG, participants listened to matrix sentences, presented at the same subject-specific SNRs as during the behavioral experiment. This way, we could link the two behavioral listening effort measures (RTprim and RTsecond; see Behavioral experiment) with envelope tracking. Since listening effort can also be measured using self-report or EEG (Pichora-Fuller *et al*., 2016), we collected three additional listening effort measures (self-report, phase synchronization and alpha power) during the EEG experiment. In the next section, the procedure of the EEG experiment, the self-report and neural effort measures as well as the signal processing using MATLAB R2016b (Mathworks, Natwick, USA) are described in more detail.

#### Neural envelope tracking and self-reported listening effort

To measure envelope tracking, we presented a concatenation of 40 matrix sentences at the same seven SNRs as the dual-task (six subject-specific SNRs and No noise). Between the sentences, a silent gap that was uniformly distributed between 0.8 and 1.2 s, was inserted. The participants listened to three repetitions of the same matrix sentences at the seven SNRs (3 repetitions × 7 SNRs = total of 21 conditions). Since matrix sentences do not contain context and are difficult to remember, repeated listening should not have introduced a bias of content learning on our results. The order of the SNRs was quasi-randomized across repetitions and participants. To keep them attentive, a question was asked per condition (e.g. “How many red boats did Jacob carry?”). After the question, participants were instructed to rate 1) their perceived understanding and 2) the effort they had to expend during listening by reporting a percentage between 0 and 100%. To better compare the five effort measures, the effort ratings were normalized per repetition similarly to the reaction times. More specifically, per participant and per SU-level, each rating was subtracted by the median for the “No noise” condition. We did not divide by the median because many participants rated the “No noise” condition equal to zero, i.e. low effort. Lastly, the story “Milan” by Stijn Vranken (8 subjects) or the story “De Wilde Zwanen” by Hans Christian Andersen (5 subjects) was presented without a masker to train the linear decoder (see Signal Processing: Envelope reconstruction).

#### Signal processing: Envelope reconstruction

Neural envelope tracking was measured by correlating the original envelope of the matrix sentences with the envelope reconstructed from the EEG responses. To obtain this, the original speech envelope was first extracted by filtering the target speech stimulus using a Gammatone filterbank followed by a power law (Biesmans *et al*., 2017). Next, the envelope was downsampled from 44100 Hz to 256 Hz to decrease processing time. Then, a type 2 zero-phase Chebyshev filter (with 80 dB attenuation at 10 % outside the passband) was applied to the envelope. For the delta-band, we filtered from 1 up to 4 Hz, for the theta-band from 4 up to 8 Hz. Finally, the speech envelope was further downsampled to 128 Hz.

The EEG data was downsampled from 8192 Hz to 256 Hz similarly to the envelope. Next, a generic EEG artifact removal algorithm (Somers *et al*., 2018) was applied on the EEG-data to remove eye artifacts similar to described in Decruy *et al*. (2019). EEG-signals were then re-referenced to Cz and bandpass filtered for delta- and theta-band using the same Chebyshev filter as used for the envelope. In a last step, the EEG was downsampled to 128 Hz.

To reconstruct the envelope from the EEG, we used the stimulus reconstruction approach described by Vanthornhout *et al*. (2018). More specifically, we used a linear decoder or spatiotemporal filter that linearly combines the EEG signals of the different channels and their time shifted versions to optimally reconstruct the envelope. Mathematically, this can be formulated as follows: *ŝ*(*t*) = ∑_*n*_ ∑_*τ*_ *g*(*n, τ*)*R*(*t* + *τ, n*), with *ŝ*(*t*) the reconstructed envelope, *t* the time ranging from 0 to T, *n* the index of the recording electrode ranging from 1 to 63, *τ* the post-stimulus integration window length, *g* the linear decoder and *R* the shifted neural responses. We chose an integration window from 0 until 250 ms. The weights of the decoder were determined in a training phase by applying ridge regression with regularization on the inverse autocorrelation matrix: *g* = (*RR*^*T*^)^*-*1^(*RS*^*T*^) with *R* as the time-lagged matrix of the EEG signal and *S* the speech envelope (inspired on the mTRF toolbox: Lalor *et al*., 2006, 2009). The decoder *g* was trained for each participant on the story “De Wilde Zwanen” or “Milan”.

After training, the subject-specific decoder was applied on the EEG responses to the matrix sentences, to reconstruct the envelope for each presentation and SNR. This reconstruced envelope was then correlated with the actual envelope using a bootstrapped Spearman correlation after removing the parts that contained silences. Neural envelope tracking per participant and per SNR condition was measured by calculating the mean of the correlations of the three presentations. A significance level for envelope tracking was calculated by correlating random permutations of the actual and reconstructed envelope (1000 times) and taking percentile 2.5 and 97.5 to obtain a 95% confidence interval.

#### Phase synchronization and alpha power

Next to RTprim, RTsecond and self-reported effort, we included two additional neural measures of listening effort in the present study. The phase synchronization of EEG responses was calculated based on the method of Bernarding *et al*. (2014). Alpha power was extracted using Thomson’s multitapers spectral analysis (Thomson, 1982). In the following section, the several signal processing steps are described in more detail.

#### Signal Processing: Phase synchronization and alpha power

The EEG was first downsampled from 8192 Hz to 512 Hz, similar to Bernarding *et al*. (2014). Then artifacts were removed using a generic EEG artifact removal algorithm (Somers *et al*., 2018). Consequently, EEG signals were re-referenced to Cz and bandpass filtered using the same Chebyshev filter as used for envelope reconstruction. In contrast to neural envelope tracking, we did not limit the filtering to the delta- or theta-band but selected a wide range of frequencies, from 1 up to 40 Hz, similar to Bernarding *et al*. (2014).

The phase synchronization was evaluated per SNR by computing the distribution of the instantaneous phase on the unit circle (Bernarding *et al*., 2014). First, we extracted the instantaneous phase per time sample (approximately 120 s of EEG-data per SNR × 512 Hz = 61440 samples) using a continuous wavelet transform (scale a = 40; Morlet wavelet). Since, we were only interested in the phase synchronization to the stimulus, we removed the instantaneous phases for the silent gaps that were inserted between the matrix sentences. Second, we evaluated the distribution of the remaining instantaneous phases for electrode TP8 (i.e., closest to the right mastoid) on the unit circle using a Rayleigh Test (Strauss *et al*., 2010; Bernarding *et al*., 2014). Therefore, we first computed the mean resultant vector of the phase values for n time samples: 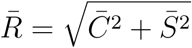 in which 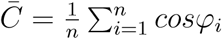 and 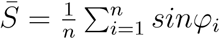. Then, we approximated the probability *Pr* that the phases are uniformly distributed using 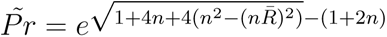. If the phases are dispersed around the unit circle, the mean resultant vector 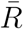 is small, resulting in a small probability to reject the null hypothesis of the test that the data are uniformly distributed. Finally, we calculated phase synchronization as 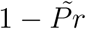. During the statistical analyses, we observed a large variability of the phase synchronization values as a function of time. Because this could be due to attentional effects in the beginning and end of each condition, we divided each condition in three equal time sequences and only included the phase values of the middle part for our next analyses.

Alpha power was calculated by extracting the power density of the EEG for frequencies between 8 and 15 Hz. We chose a broad frequency range because alpha power can peak at a different frequencies between participants. Per SNR condition, power density for this frequency range was extracted from the EEG measured during presentation of matrix sentences. This was done over consecutive non-overlapping 1-s epochs, using Thomson Multitapers Spectral Analysis (7 tapers; Thomson, 1982). Topoplots on which the median of alpha power across epochs was plotted, showed only changes across SU-levels for the parietal-occipital electrodes. Because of this, we selected a subset of electrodes (P9, P7, P5, PO3, POz, PO4, P6, P8, P10, PO7, O1, O2, Oz, PO8, Iz) and used the median alpha power across this selection and time course as our second neural effort measure.

Finally, phase synchronization as well as alpha power were normalized per repetition, SNR and participant. In previous studies, alpha power was often normalized against a baseline period before the stimulation (Obleser *et al*., 2012; Dimitrijevic *et al*., 2019). We did not do this because we wanted to compare the five effort measures, so we normalized each measure the same way, i.e. each value was subtracted and divided by the median of the “No noise” condition.

### EEG recording

The EEG experiment was administered for all participants at the research group in a triple-walled, soundproof booth, equipped with a Faraday cage to avoid electromagnetic interference. The same auditory stimulation set-up as the behavioral experiment, was used to present the matrix sentences to the participants. To measure the EEG, we mounted 64 active Ag/AgCl electrodes and two extra electrodes, serving as the common electrode (CMS) and current return path (DRL), in head caps according to the 10-20 electrode system. The EEG recordings were conducted using the BioSemi ActiveTwo system (Amsterdam, Netherlands), digitized at a sampling rate of 8192 Hz and stored on a hard disk using the BioSemi ActiView software.

### Statistical analyses

The statistical analyses were conducted using R software (version 3.4.4; nlme package - version 3.1-131.1; Field et al., 2012; Pinheiro *et al*., 2017). We investigated the association between the different listening effort measures and their link with envelope tracking using Linear Mixed-effect Models (LMMs). The fixed-effect part of the LMMs consisted of the predictors of interest whereas the random-effect part included the variable participant, nested in the repeated measure predictor SU-level if this improved the model fit. All models were fitted using the maximum likelihood estimation. The best fitting model was determined by first progressively introducing multiple fixed-effects and corresponding interactions. Then, the different models were compared using likelihood ratio tests and Akaike’s Information Criterion (AIC; Akaike, 1974). The best fitting model served as starting point for the evaluation of the contribution of other predictors until the final best fitted model was determined. Significant main and interaction effects of the final model are discussed in the results section by reporting the unstandardized regression coefficient (*β*) with standard error (SE), degrees of freedom (df), t-Ratio and p-value per fixed-effect term. A significance level of *α* = 0.05 was set for all the models.

## Results

### Listening effort as a function of speech understanding

Prior to linking each effort measure with neural envelope tracking, we first compared the different listening effort measures based on how they behave as a function of speech understanding.

#### Speech-in-noise performance

The five listening effort measures used in this study were collected during different experiments. RTprim and RTsecond were collected during the behavioral experiment whereas self-report, phase synchronization and alpha power were registered during the EEG experiment. Although we used the same subject-specific SNRs for each experiment, differences in speech understanding performance could result in differences between the measures. We investigated this by comparing the speech understanding scores obtained during three procedures: the recall scores during the speech-in-noise test administered as a (1) single-task paradigm (2) or dual-task paradigm as well as (3) the rated scores obtained during the EEG experiment. The psychometric functions, plotted in figure 3, show that similar speech understanding scores were obtained for the three procedures. Accordingly, no significant effect was detected by an LMM with procedure as fixed-effect term and participant nested in SU-level as random-effect term (p > 0.05). Therefore, we will not make a distinction between the recall versus rated speech understanding scores for the following analyses.

**Figure 3:**
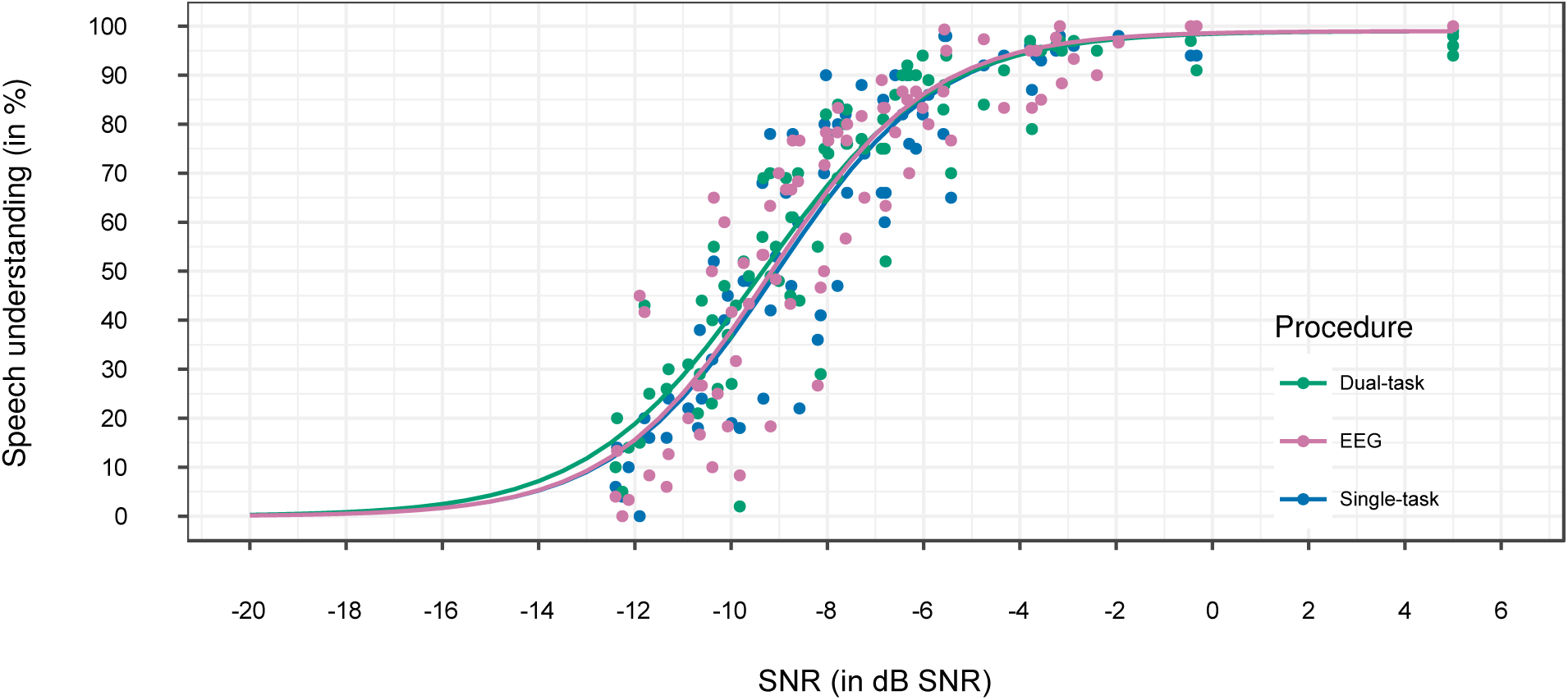
Comparison of speech in noise performance of 13 participants measured during the single-task paradigm (recall), dual-task paradigm (recall) and EEG experiment (rate). For each procedure, a psychometric function (color-coded) is fitted on the data of all participants across the six subject-specific SNRs and No noise condition (for visualization No noise condition is plotted at 5 dB SNR instead of 100 dB SNR). No significant difference was found between the different procedures.

#### Behavioral, self-report and neural measures of listening effort

To compare the five listening effort measures, we analyzed each outcome measure as a function of the speech understanding scores obtained at the seven SNRs (six subject-specific SNRs and No noise). Per measure, we built an LMM with speech understanding as fixed-effect term (table 1; figure 4). For the behavioral experiment, we detected a supralinear decrease in RTprim with increasing speech understanding (*β* = −2.51e-04, SE = 3.88e-05, p < 0.001). Moreover, RTprim was long, i.e. worse, at unfavorable SU-levels (0-30%) and then became shorter, i.e. better, with increasing speech understanding (see figure 4, left panel). RTsecond, on the other hand, did not decrease as a function of speech understanding, but had a peak-shape (revealed by a significant cubic and linear fixed-effect term; cubic: *β* = −9.22e-07, SE = 2.53e-07, p < 0.001; linear: *β* = 7.31e-03, SE = 3.23e-03, p = 0.026). This can also be inferred from figure 4 in which RTsecond first increased from unfavorable SU-levels (0-10%) until intermediate SU-levels (40-60%), and then decreased with increasing speech understanding (80-100%). Taken together, RTprim and RTsecond showed a different effect of listening effort as function of speech understanding.

**Table 1:** Linear Mixed-effect Models: The effect of speech understanding on five listening effort measures (RTprim, RTsecond, Self-report, Phase synchronization and Alpha power). Regression coefficients (*β* values), standard errors (SE), degrees of freedom (df), t-Ratios and p-values are reported per fixed-effect term. Participant was included as a random effect.

| Behavioral experiment |  |  |  |  |  |  |
| --- | --- | --- | --- | --- | --- | --- |
| Measure | Fixed-effect terms | $\beta$ value | SE | df | t-Ratio | p-value |
| RTprim | Intercept | 2.41 | 0.38 | 77 | 6.33 | < 0.001 |
|  | SU<sup>2</sup> | -2.51e-04 | 3.88e-05 | 77 | -6.47 | < 0.001 |
| RTsecond | Intercept | 0.20 | 0.13 | 76 | 1.58 | 0.117 |
|  | SU | 7.31e-03 | 3.23e-03 | 76 | 2.27 | 0.026 |
|  | SU<sup>3</sup> | -9.22e-07 | 2.53e-07 | 76 | -3.65 | < 0.001 |
| EEG experiment |  |  |  |  |  |  |
| Measure | Fixed-effect terms | $\beta$ value | SE | df | t-Ratio | p-value |
| Self-report | Intercept | 92.11 | 3.25 | 77 | 28.34 | < 0.001 |
|  | SU<sup>2</sup> | -8.62e-03 | 2.83e-04 | 77 | -30.41 | < 0.001 |
| Phase | Intercept | 6.69 | 2.87 | 77 | 2.33 | 0.023 |
|  | SI | -0.03 | 0.03 | 77 | -1.00 | 0.318 |
| Alpha | Intercept | 0.05 | 0.06 | 76 | 0.87 | 0.385 |
|  | SU<sup>2</sup> | 9.31e-05 | 3.20e-05 | 76 | 2.91 | 0.005 |
|  | SU<sup>3</sup> | -9.32e-07 | 3.06e-07 | 76 | -3.04 | 0.003 |

**Figure 4:**
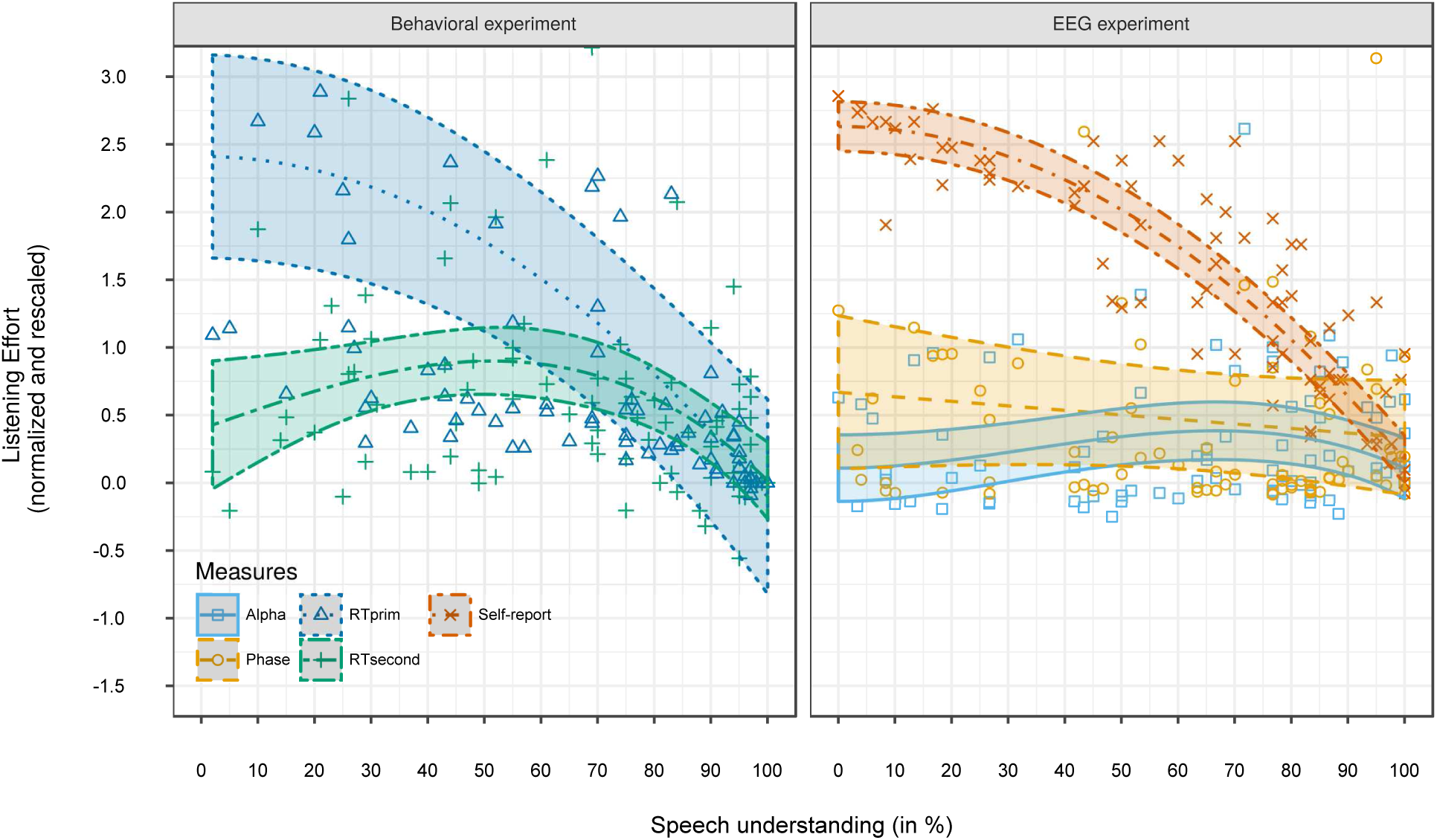
Listening effort as a function of speech understanding for two measures obtained during the behavioral experiment (RTprim and RTsecond, left panel) and for three measures obtained during the EEG experiment (self-report, phase synchronization and alpha power, right panel). For each measure, a regression line with confidence intervals (shaded areas; color-coded) was fitted on the data. The regression lines indicate that different measures reflect different aspects of listening effort (see table 1).

From the EEG experiment, we collected three listening effort measures. Similar to RTprim, self-report decreased supralinearly with increasing speech understanding (*β* = −8.62e-03, SE = 2.83e-04, p < 0.001, see also right panel of figure 4). The neural measures of listening effort behaved differently. More specifically, the LMM for phase synchronization detected no significant effect of speech understanding (p > 0.05). The LMM for alpha detected a peak-shape as a function of speech understanding (revealed by a significant cubic and quadratic fixed-effect term; cubic: *β* = −9.32e-07, SE = 3.06e-07, p = 0.003; quadratic: *β* = 9.31e-05, SE = 3.20e-05, p = 0.005; figure 4). More specifically, alpha power was highest between 60% and 80% speech understanding.

### Neural envelope tracking and listening effort

The second aim of this study was to investigate how listening effort and neural envelope tracking are associated. First, we studied the effect of speech understanding on neural envelope tracking. Second, we analyzed which listening effort measures contribute to the unexplained variance in neural envelope tracking.

#### Neural envelope tracking

To investigate the association between speech understanding and neural envelope tracking, we built an LMM with speech understanding as fixed-effect term and participant nested in frequency band as random-effect. The LMM revealed that envelope tracking was best predicted, i.e. lowest AIC, using speech understanding as a cubic fixed-effect term (*β* = 5.51e-08, SE = 6.39e-09, p < 0.001). As shown in figure 5, envelope tracking increased with speech understanding. In addition to speech understanding, the LMM detected a significant main effect of frequency band. More specifically, lower envelope tracking was observed for theta compared to delta (*β* = −0.02, SE = 7.21e-03, p = 0.011). No significant interaction was observed between speech understanding and frequency band.

**Figure 5:**
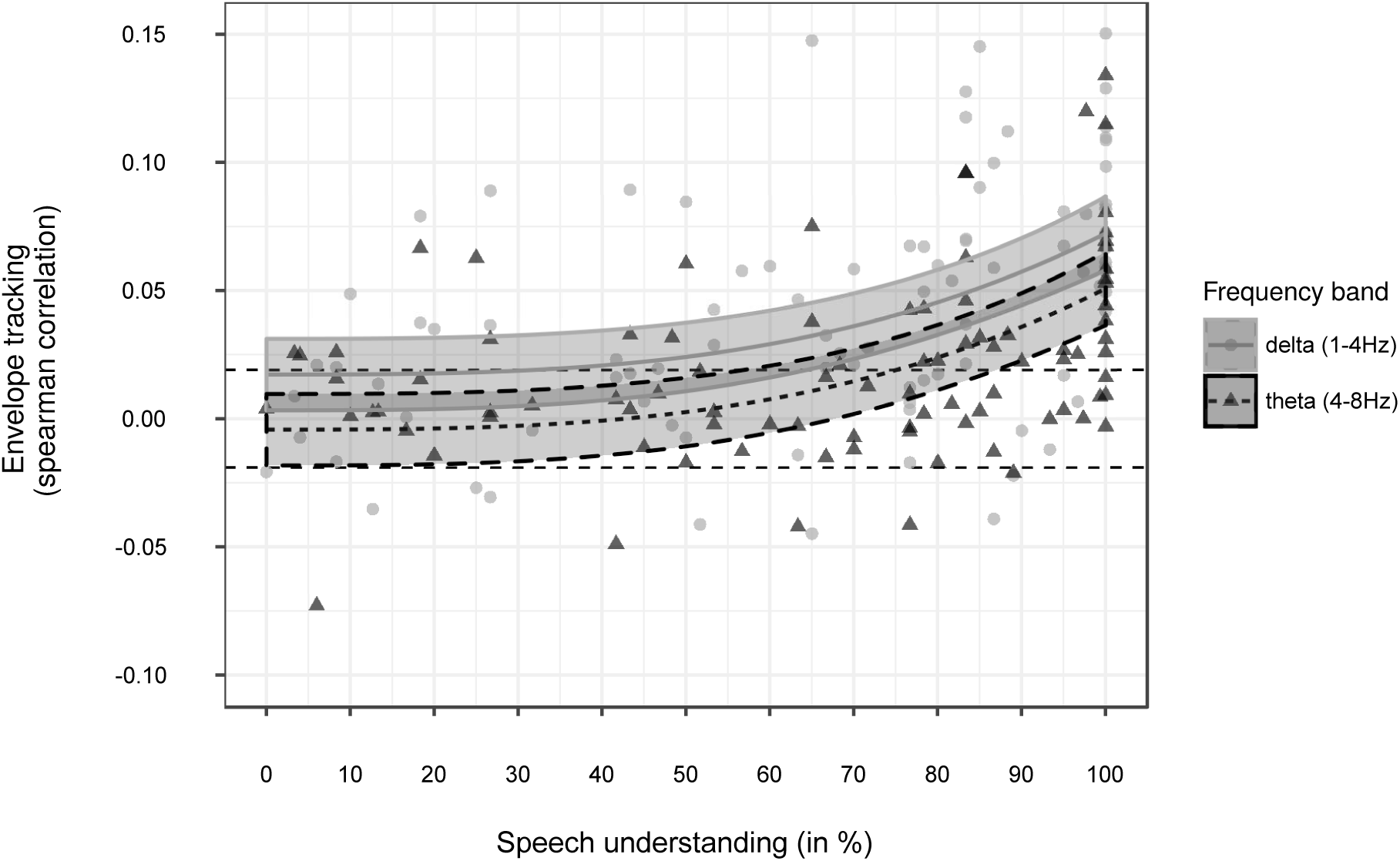
Neural envelope tracking as a function of speech understanding, measured in 13 participants. The regression lines with confidence intervals (shaded areas) indicate that envelope tracking increases supralinearly with increasing speech understanding for both frequency bands. In addition, lower envelope tracking was obtained for theta (4-8 Hz) versus delta (1-4 Hz) band (color-coded). Dashed black lines indicate the significance level of the measure for envelope tracking.

### Link between listening effort and envelope tracking

We assessed per frequency band whether listening effort accounts for a part of the inter-subject variability in envelope tracking by adding each listening effort measure to the basis model (with SU^3^ as predictor for envelope tracking). Only for the theta-band (4-8 Hz), adding two of the five measures resulted in a significant improvement of the model fit. Likelihood ratio tests and the AIC showed no improvement for RTprim, alpha or phase. For RTsecond and self-report, on the other hand, a better model fit was achieved when adding these measures as an interaction with SU^3^. For both RTsecond and self-report, envelope tracking decreased with increasing effort when participants understood the sentences well whereas envelope tracking increased with increasing effort when it was difficult to understand speech (RTsecond: *β* = −8.38e-08, SE = 2.91e-08, p = 0.005; self-report: *β* = −8.15e-10, SE = 3.54e-10, p = 0.024; see table 2). Although adding these effort measures resulted in a significantly better model fit, we only found an improvement in Pseudo-R^2^ of 0.054 when adding RTsecond and 0.043 when adding self-reported effort (Pseudo-R^2^ for basis model: 0.27). Taken this into account, the effect of listening effort on neural envelope tracking is rather small.

**Table 2:** Linear Mixed-effect Models: The effect of listening effort on neural envelope tracking. Regression coefficients (*β* values), standard errors (SE), degrees of freedom (df), t-Ratios and p-values are reported per fixed-effect term. Participant was included as a random effect.

| Measure | Fixed-effect terms | $\beta$ value | SE | df | t-Ratio | p-value |
| --- | --- | --- | --- | --- | --- | --- |
| RTsecond | Intercept | 1.27e-04 | 7.53e-03 | 75 | 0.02 | 0.987 |
|  | SU<sup>3</sup> | 5.33e-08 | 1.07e-08 | 75 | 4.98 | < 0.001 |
|  | RTsecond | 0.01 | 0.01 | 75 | 0.96 | 0.342 |
|  | RTsecond:SU<sup>3</sup> | -8.38e-08 | 2.91e-08 | 75 | -2.88 | 0.005 |
| Self-report | Intercept | 0.01 | 0.02 | 75 | 0.62 | 0.540 |
|  | SU<sup>3</sup> | 3.96e-08 | 2.56e-08 | 75 | 1.54 | 0.127 |
|  | LEsubj | -7.15e-05 | 2.90e-04 | 75 | -0.25 | 0.806 |
|  | LEsubj:SU<sup>3</sup> | -8.15e-10 | 3.54e-10 | 75 | -2.30 | 0.024 |

## Discussion

The aim of this study was to investigate if listening effort modulates neural envelope tracking. First, we studied the behavior of five effort measures as a function of speech understanding. We found different outcomes among the different measures. Self-report and reaction times on the speech-in-noise test showed a consistent decrease in effort with increasing speech understanding. Alpha power and reaction times on the verbal working memory test, on the other hand, showed maximum effort at the intermediate instead of lowest speech understanding levels. Second, our results revealed a significant increase in neural envelope tracking with increasing speech understanding. Only two of the five effort measures accounted for a small part of the inter-subject variability in envelope tracking, suggesting that listening effort does not substantially modulate envelope tracking in young, NH listeners.

### Different measures reflect different aspects of listening effort

#### Behavioral, self-report and neural measures of listening effort

To compare the five listening effort measures, we analyzed each measure as a function of speech understanding (figure 4). Our results show that the self-report measure and RTprim both quadratically decrease with increasing speech understanding. In other words, highest effort was observed at unfavorable SU-levels whereas lowest effort was obtained when participants understand speech best. This behavior is in line with the findings of previous studies (Gatehouse & Gordon, 1990; Luts *et al*., 2010; Zekveld *et al*., 2010; Houben *et al*., 2013; Wendt *et al*., 2014). For example, Rudner *et al*. (2012) asked hearing impaired participants to rate their listening effort using a visual analog scale, across five SU-levels (between 50% SU and 95% SU). Their results showed that perceived effort consistently decreased from the highest to lowest SU-level. Similarly, Pals *et al*. (2015) found longer RTprim in the most difficult conditions and shortest RTprim in the “No noise” condition.

In contrast to this, we found that RTsecond and alpha power were both best modeled using a cubic term, resulting in a peak-shaped behavior. More specifically, RTsecond was short, i.e. low effort, at the extreme unfavorable or favorable SU-levels whereas RTsecond was long at intermediate SU-levels (40-60%). These results are in agreement with the findings of dual-task studies (Wu *et al*., 2016) as well as pupillometry studies (Ohlenforst, Zekveld, & Lunner *et al*., 2017) that investigated listening effort across a wide range of SNRs. Similarly, highest alpha power was obtained at intermediate SU-levels (60-80%). Although increased alpha power has been associated with increased demands or effort (Obleser *et al*., 2012; Dimitrijevic *et al*., 2019), to our knowledge no research has shown a peak-shaped behavior for alpha power similar to the present study.

In sum, the consistent decrease in effort versus peak-shape indicate that different methods indeed tap into different aspects of listening effort (Lemke & Besser, 2016; Alhanbali *et al*., 2019).

#### Perceived effort versus processing load

Since self-reported effort and RTprim behave similarly in the present study, we could assume that they reflect the same aspect of listening effort. However, self-report measures are often considered to measure perceived effort whereas single-task paradigm reaction times (RTprim) are assumed to reflect the cognitive processing time required to understand speech (Pals *et al*., 2015). A possible explanation could be that these measures behave similarly because in our study self-report and RTprim did not measure the motivation of the participant. In the FUEL framework of Pichora-Fuller *et al*. (2016), listening effort is modeled as a function of both demands as well as the motivation of a person to complete the task. Self-reported effort was measured by the question “How much effort did you have to expend to understand the sentences?”. This is somewhat ambiguous, because participants could interpret this question in different ways. Recent studies have suggested that participants tend to substitute effort questions by an easier, perceived performance question (Moore & Picou, 2018; Picou & Ricketts, 2018), resulting in highest effort ratings for the most unfavorable SNRs. This could be solved by asking questions without explicitly using the term “effort” (Moore & Picou, 2018). For example, the question “How hard were you trying to understand the speech” (Wu *et al*., 2016) or asking to also indicate how often they had given up trying to perceive the sentences (Zekveld *et al*., 2014), can both give an indication of the aspect motivation and could consequently have resulted in a peak-shaped curve similar to RTsecond (Wu *et al*., 2016). When examining listening effort at a single SNR, Picou & Ricketts (2018) advise to use ratings about the desire to improve the listening condition in order to measure expended effort. Similar to the self-report effort questions, we believe that RTprim can also be biased by the performance on the speech-in-noise test whereas dual-task performance (RTsecond) solely measures the performance on the secondary task. When participants give up, automatically more resources will be available resulting in a better performance on the secondary task. For RTprim on the other hand, bad performance on the speech-in-noise test can result in a longer processing time to decide what to answer.

No consensus is yet established about which listening effort aspect alpha power and phase synchronization represent. Our results have shown no effect of speech understanding on phase synchronization. There are a few methodological differences with Bernarding *et al*. (2014) that could explain this. For example, their participants were instructed to recall sentences whereas our participants needed to listen to concatenated sentences. When listening to sentences instead of recalling, attention can drift away more easily resulting in a weaker phase synchronization. This however does not seem a likely reason as we asked questions to keep participants alert. Secondly, Bernarding *et al*. (2014) varied effort by changing hearing aid settings while we manipulated effort by changing speech understanding. In view of this, several factors seem to affect the sensitivity of phase synchronization to be used as a robust neural correlate of listening effort. In contrast to phase, our results showed a peak-shaped curve for alpha power as a function of speech understanding. Although, no study has yet investigated changes in alpha power across a wide range of SNRs or SU-levels, several studies demonstrated that increased alpha is observed with increased demand or effort (Obleser *et al*., 2012; Miles *et al*., 2017; Dimitrijevic *et al*., 2019).

### Neural envelope tracking and listening effort

#### Neural envelope tracking as a function of speech understanding

We measured neural envelope tracking for two frequency bands. For both the delta- and theta-band, an increase in neural envelope tracking with increasing speech understanding was observed. This is in line with the findings of previous studies in which neural envelope tracking of intelligible sentences or story was measured as a function of SNR (Vanthornhout *et al*., 2018; Etard & Reichenbach, 2019; Lesenfants *et al*., 2019). Hence, our results further confirm the potential of neural envelope tracking as an objective measure of speech understanding (Vanthornhout *et al*., 2018; Lesenfants *et al*., 2019).

#### Listening effort does not substantially modulate neural envelope tracking

Recent research has demonstrated the influence of several top-down processes on neural envelope tracking. We investigated whether the effort that participants expend when listening to matrix sentences modulates envelope tracking. For the delta-band, no association was found between listening effort and neural envelope tracking. These results are in line with a recent study of Müller *et al*. (2019) in which listening effort was measured using self-report and pupillometry, and neural envelope tracking using the cross-correlation between the onset envelope and EEG response. Even though all measures were similarly affected by speech rate, no significant correlations were found between the listening effort and neural envelope tracking. According to the authors, this is due to different factors influencing neural envelope tracking versus self-reported effort and pupillometry.

For the theta-band, two of our five listening effort measures, RTsecond and self-reported effort, explained a significant part of the inter-subject variability of neural envelope tracking. This is in agreement with the results of Wisniewski *et al*. (2015), showing a greater power for frontal midline theta-band activity when participants expend more effort during a speech recognition task. In spite of this, we have to note that only a small part of the inter-subject variability in neural envelope tracking was explained by our listening effort measures. Taking this into account, our results suggest that neural envelope tracking is not substantially modulated by listening effort.

Lastly, the observed small effect of listening effort on neural envelope tracking was not completely in line with our original hypothesis. Based on previous studies, we expected to find lower envelope tracking for speech without a masker compared to speech presented at a high SNR due to lower effort. For example, Das *et al*. (2018) found lower neural envelope tracking for stories presented without a masker compared to stories presented at −1.1 dB SNR. Similarly, Lesenfants *et al*. (2019) observed lower envelope tracking in the theta-band for matrix sentences presented without a masker compared to sentences presented at −3.5, −1, −0.5 or 2.5 dB SNR. As shown in figure 5, we found, in contrast to these studies, highest envelope tracking in the “No noise” condition in which 100% speech understanding was achieved. Therefore, adding our listening effort measures to the basis model for neural envelope tracking could not have resulted in the expected effect. More research is needed to unravel the observed lower envelope tracking to sentences presented without a masker in previous studies.

### Future work

Recently, studies have shown that envelope tracking increases with advancing age. Older NH adults showed an enhanced envelope tracking, which appears to be associated with the age-related deterioration of cognitive processes (Presacco *et al*., 2016; Decruy *et al*., 2019). Taking into account that older adults expend more effort during listening (Degeest *et al*., 2015), this enhanced envelope tracking could reflect increased listening effort. Although we have only found a small effect in the present study, including effort measures in future aging studies could be beneficial to further investigate this. Based on our results in young NH participants, we would suggest to use self-report measures to quantify listening effort as they can be very easily implemented in an existing speech-in-noise test. In addition, studies have shown that self-reported effort can not only reflect perceived effort but also processing load if participants are also instructed to indicate when they give up (Zekveld *et al*., 2014; Wu *et al*., 2016). This way, demands as well as motivation, which are suggested to both affect effort, will be measured (Pichora-Fuller *et al*., 2016; Peelle, 2018). However, we have to note that older adults tend to underestimate their degree of difficulties (Anderson Gosselin & Gagné, 2011). As this can affect self-reported effort, it should be investigated whether alpha power or behavioral effort measures correlate better with the enhanced envelope tracking in older NH adults.

## Conclusion

In the present study, we investigated listening effort across a wide range of subject-specific SNRs. Our results reveal that different measures indeed lead to different outcomes and support the importance of including a measure of motivation as well as demands to reliably quantify listening effort. In line with previous studies, we observed an increase in envelope tracking with increasing speech understanding. Since no substantial influence of listening effort was found on envelope tracking for our participants, our results suggest that effort may not confound the potential of neural envelope tracking to objectively measure speech understanding.

## Acknowledgments

The authors want to thank all participants for their participation in this study as well as research student Maud Lobel for her help in participant recruitment and data acquisition.

## Author Contributions

L.D., D.L., J.V., T.F. conceived and designed the research; L.D. performed experiments;

L.D. analyzed data; L.D., D.L., J.V., T.F. interpreted results of experiments; L.D. prepared figures; L.D, D.L., J.V., T.F. drafted the manuscript.

## Abbreviations

dB SNR: decibel Signal-to-noise ratio
dB HL: decibel hearing level
dB SPL: decibel sound pressure level
df: degrees of freedom
EEG: electroencephalography
LMM: Linear Mixed-effect Model
MAD: median absolute deviation
NH: normal-hearing
RTprim: Reaction time on primary task
RTsecond: Reaction time on secondary task
SE: standard error
SNR: signal-to-noise ratio
SRT: speech reception threshold
SU: speech understanding
SWN: speech-weighted noise

